# Structural flexibility of apolipoprotein E-derived arginine-rich peptides improves their cell penetration capability

**DOI:** 10.1101/2023.08.16.553537

**Authors:** Yuki Takechi-Haraya, Takashi Ohgita, Akiko Usui, Kazuchika Nishitsuji, Kenji Uchimura, Yasuhiro Abe, Ryuji Kawano, Monika I. Konaklieva, Mart Reimund, Alan T. Remaley, Yoji Sato, Ken-ichi Izutsu, Hiroyuki Saito

## Abstract

Amphipathic arginine-rich peptide, A2-17, exhibits moderate perturbation of lipid membranes and the highest cell penetration among its structural isomers. We investigated the direct cell-membrane penetration mechanism of the A2-17 peptide. We designed structurally constrained versions of A2-17, stapled (StpA2-17) and stitched (StchA2-17), whose α-helical conformations were stabilized by chemical crosslinking. Circular dichroism confirmed that StpA2-17 and StchA2-17 had higher α-helix content than A2-17 in aqueous solution. Upon liposome binding, only A2-17 exhibited a coil-to-helix transition. Confocal microscopy revealed that A2-17 had higher cell penetration efficiency than StpA2-17 in HeLa cells. Partitioning into lipid membranes was more prominent for StchA2-17 than for A2-17 or StpA2-17; StchA2-17 remained on the cell membrane without cell penetration. Tryptophan fluorescence analysis suggested that A2-17 and its analogs had similar membrane-insertion positions between the interface and hydrophobic core. Atomic force microscopy demonstrated that A2-17 reduced the mechanical rigidity of liposomes to a greater extent than StpA2-17 and StchA2-17. Finally, electrophysiological analysis showed that A2-17 induced a higher charge influx through transient pores in a planer lipid bilayer than StpA2-17 and StchA2-17. These findings indicate that structural flexibility, which enables diverse conformations of A2-17, leads to a membrane perturbation mode that contributes to cell membrane penetration.

## Introduction

In recent years, there have been substantial advances in the development of various drug modalities as active pharmaceutical ingredients (APIs), such as low molecular weight compounds, physiological peptides/proteins, and functional nucleotides^1-3^. However, the clinical use of these APIs is hindered by their limited cell penetration or membrane permeation capabilities^4^. Arginine-rich peptides (ARPs) have garnered considerable attention as they can efficiently deliver different APIs inside cells, both in vitro and in vivo^5,6^. Understanding the entry mechanism of ARPs into cells is crucial for developing targeted therapeutics and for establishing a universal methodology to design, evaluate, and ensure the efficacy and safety of substances of interest.

The endocytic uptake pathway is likely the primary cell entry mechanism of ARPs, when delivering APIs with large molecular weight, such as proteins and nucleotides^5-7^. However, certain types of ARPs, such as polyarginine, Tat, Rev, and A2-17, when conjugated with or without low molecular weight compounds, demonstrate non-endocytic entry predominance in cells at 4 °C, where all the membrane trafficking mechanisms, including endocytosis, are suppressed^8-10^. This direct cell membrane penetration process is a spontaneous physicochemical phenomenon independent of specific membrane proteins or energy derived from the hydrolysis of adenosine triphosphate^10-12^. The direct penetration process likely occurs by a membrane perturbation mechanism, wherein the interaction between Arg residues of peptides and oxonium anion groups, including phosphate groups of lipids, perturbs and induces the lipid bilayer to locally form short-life transient non-lamellar defects or pores; which then enables ARPs to traverse the membrane barrier^12-14^.

Amphipathic helical structures can be used to develop highly cell-penetrable ARPs, as observed in many cell-penetrating peptides^6,12,15^. The amphipathic ARP peptide called A2-17 was designed based on the glyocosaminoglycan-binding region of human apolipoprotein E^9,16,17^, a structural protein found on lipoproteins. It exhibits direct cell membrane penetration even at low peptide concentrations, unlike conventional ARPs, such as Tat, polyarginine, and Rev^9,10^. The increased amphipathicity of A2-17 leads to enhanced insertion and membrane perturbation, which results in generation of transient defects or pores in the membrane^18^. While the importance of conformation-restricting intramolecular linking in improving peptide cell penetration has been reported, the significance of structural flexibility in cell-penetrating peptides remains unclear^19-25^.

The present study aimed to investigate the direct cell membrane penetration mechanism of the A2-17 peptide. For this, two derivatives of A2-17, namely stapled A2-17 (StpA2-17) and stitched A2-17 (StchA2-17), were designed (Fig. 1). Their amphipathic helical structures were respectively stabilized through “hydrocarbon-stapling” with two crosslinking points and “hydrocarbon-stitching” with three crosslinking points via olefin metathesis using (*S*)-α-methyl,α-pentenylglycine (S5), (*S*)-α-methyl,α-octenylglycine (S8), and 2-(((9H-Fluoren-9-yl)methoxy)-carbonylamino)-2-(pent-4-enyl)hept-6-enoic acid (B5)^20,26^. We used a series of biophysical methods to compare the lipid membrane interaction and cell membrane penetration of these peptides.

**Figure 1.**
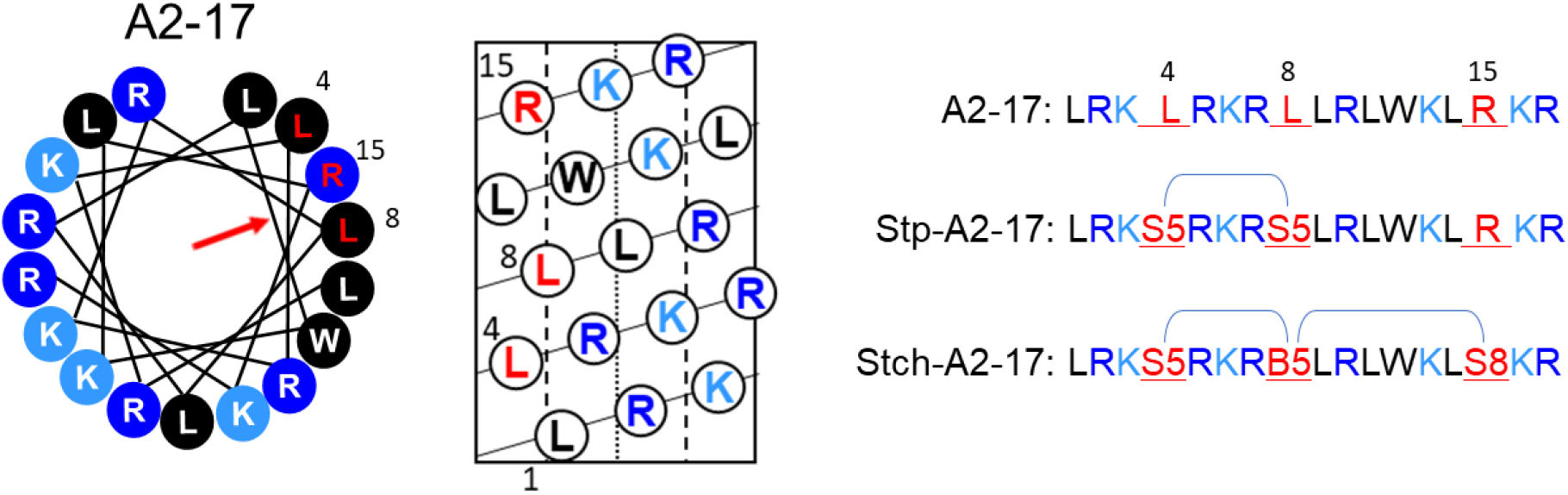
Design of stapled A2-17 (StpA2-17) and stitched A2-17 (StchA2-17) peptides. Helical wheel and helical net diagrams of A2-17 are used to display the site where the chemical linkage is introduced to StpA2-17 and StchA2-17. The helical wheel is arranged as an ideal α-helix (100° rotation per residue) as observed from the top of the long axis from the amino-terminal end. The position of red arrow in the helical wheel plot indicates the orientation of the hydrophobic moment of the peptide.

## Results

### Secondary structure of A2-17 derivatives

Circular dichroism (CD) spectra were recorded to analyze the secondary structures of intramolecularly cross-linked StpA2-17 and StchA2-17 (Fig. 2). The fourth and eighth leucine residues of A2-17 in StpA2-17 are replaced with S5. In StchA2-17, the 4th, 8th, and 15th arginine residues of A2-17 are replaced with S5, B5, and S8, respectively (Fig. 1). In water, the secondary structure of A2-17 exhibits a random coil-like conformation, as indicated by a peak around 200 nm. However, upon binding to large unilamellar vesicles (LUVs) of the cell membrane model, which was composed of distearoylphosphatidylcholine (DSPC) and distearoylphosphatidylglycerol (DSPG), it is structurally transformed into an α-helix, as revealed by double negative peaks around 206 nm and 222 nm (Fig. 2a). This behavior was similar to that observed in our previous study using small unilamellar vesicles that had a high membrane curvature^18^. In A2-17, the α-helix content in water (19%) increased by 13% upon binding to lipid membranes, although no statistically significant difference was detected (Fig. 2b). In contrast, StpA2-17 and StchA2-17 showed a distinctive CD spectrum of the α-helix structure even in water (Fig. 2a), and the calculated α-helix contents of StpA2-17 (51%) and StchA2-17 (102%) were significantly higher than that of A2-17. This suggests that intramolecular cross-linking can induce a conformational restraint (Fig. 2b). Additionally, the presence of LUVs caused no prominent secondary conformational transitions in StpA2-17 and StchA2-17, indicating that they had a stable α-helical-like conformation.

**Figure 2.**
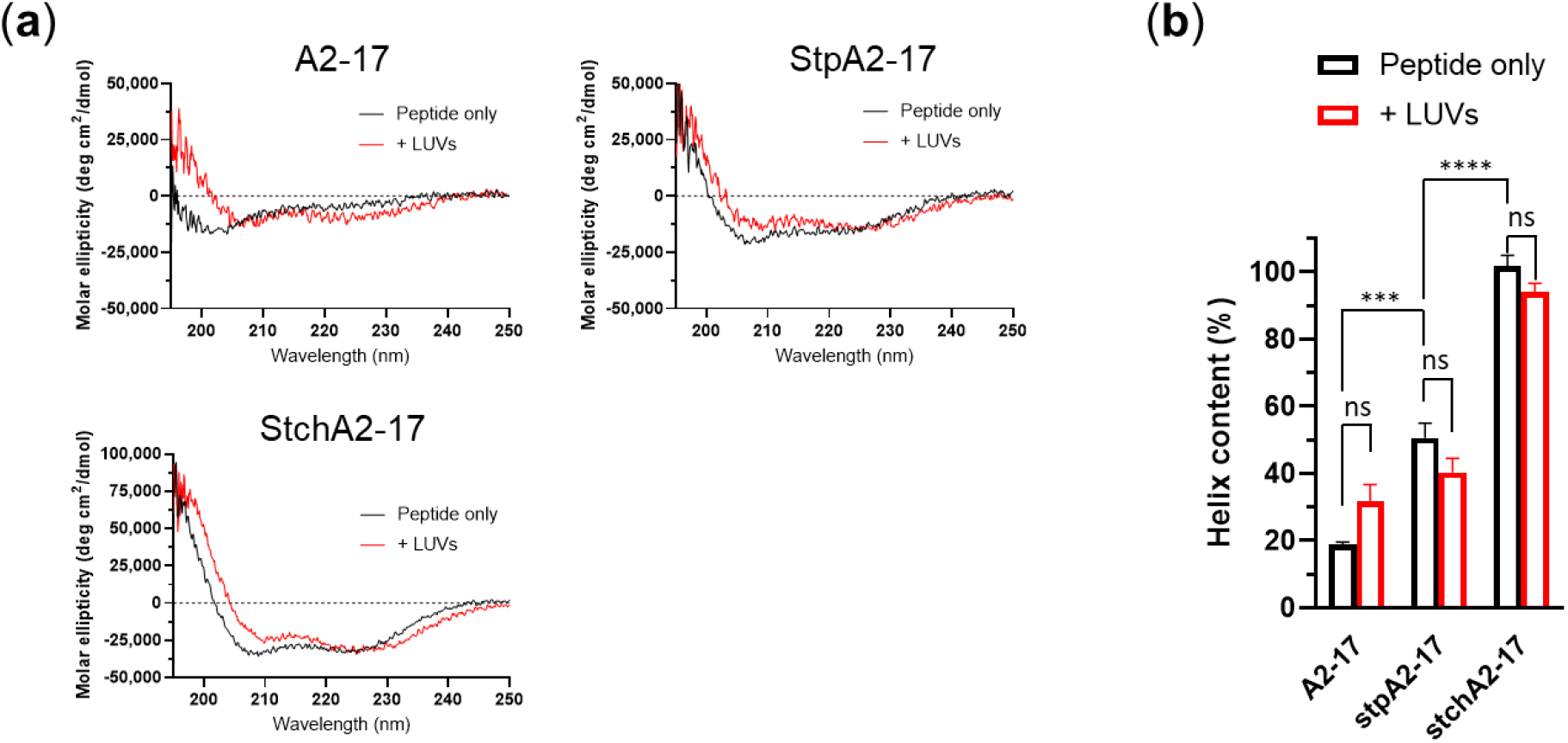
Secondary structure of peptides. (a) Far-UV CD spectra of A2-17, StpA2-17, and StchA2-17 in the absence (peptide only) and presence of lipid vesicles (+LUVs; lipid/peptide molar ratio = 100). (b) α-helix content of the peptides in the absence (peptide only) or presence of lipid vesicles (+ LUVs). \*\*\**p* < 0.001; \*\*\*\**p* < 0.0001; ns: not significant.

### Cell membrane penetration

The direct cell membrane penetration of peptides at 4 °C were compared using fluorescence detection. For this, HeLa cells treated with peptide conjugated with a small molecule drug model, 5-carboxyfluorecein (FAM), were observed using confocal fluorescent laser microscopy (Fig. 3). The intracellular FAM fluorescence intensity was in the order A2-17 > StpA2-17 > StchA-17 (Fig. 3a), and StchA2-17 remained in the peripheral region of the cells. A comparison of FAM fluorescence intensity in the cell nucleus region subtracted by extracellular FAM fluorescence intensity showed that StchA2-17 was not significantly different from the control group, which did not contain FAM-labeled peptide (Fig. 3b). These results indicated that A2-17 exhibits a higher cell membrane penetration ability than StpA2-17, and StchA2-17 could not penetrate cells.

**Figure 3.**
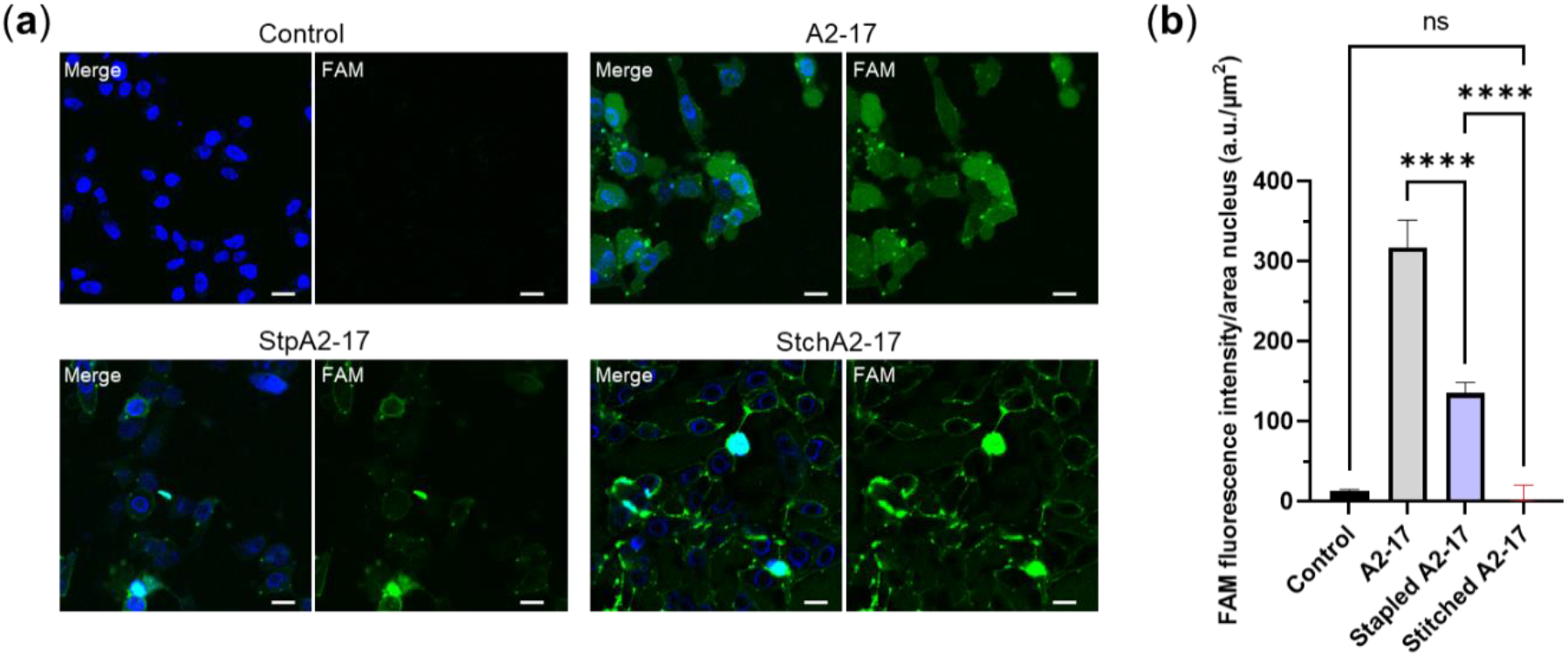
Cell membrane penetration of peptides. (a) Representative confocal fluorescent images of HeLa cells treated with 1 μM of FAM-labeled A2-17, StpA2-17, or StchA2-17 for 30 min at 4 °C. FAM fluorescence (green) and Hoechst fluorescence (blue) of counterstained nuclei are shown in the merge images (labelled as Merge in the photo) along with the FAM fluorescence images (labelled as FAM in the photo). The scale bars represent 20 μm. (b) Quantification of the FAM fluorescence intensity attributable to the peptide remaining in the nucleus region of cells. \*\*\*\**p* < 0.0001; ns: not significant.

### Lipid membrane binding

The lipid membrane-binding properties of A2-17, StpA2-17, and StchA2-17 were analyzed using their intrinsic Trp fluorescence (Fig. 4). We compared the degree of membrane insertion by monitoring the fluorescence wavelength maximum (WMF). WMF shifts toward a shorter wavelength as the peptide is transferred to a more hydrophobic environment upon binding to lipid membranes^27^. When titrated with LUVs, the WMF shift occurred faster for StchA2-17 than for A2-17 and StpA2-17 (Fig. 4a), indicating that StchA2-17 has a higher lipid binding affinity than A2-17 and StpA2-17. However, there were no significant differences between the peptides in their WMF values at the plateau when there was a large excess of LUVs over the peptides. At these conditions, only the Trp residue environment of the peptide bound to the membrane is reflected in the WMF. The WMF profiles of *N*-acetyl-L-tryptophanamide, which depend on dielectric constant (*ε*_r_), were obtained by fluorescence measurements using ethanol (see Methods for details). The Trp position of the peptide inserted into the lipid membranes was estimated from the WMF value at the plateau (Fig. 4b). All plateau WMF values were around *ε*_r_ of 20, suggesting that each peptide inserts into membranes to a similar extent between the lipid membrane interface region (*ε*_r_∼30–40) and the hydrophobic core (*ε*_r_∼2)^28,29^. The analysis of partition constant of peptides to lipid membranes using the ultrafiltration method (see Methods section for details) revealed that partition constant for StchA2-17 was approximately 50-fold larger than for A2-17 and StpA2-17 peptides, resulting in a significant increase in the standard Gibbs free energy of transfer of peptides from water to the membrane for StchA2-17 compared with A2-17 and StpA2-17 (Table 1: StchA2-17 vs A2-17, *p* < 0.001; A2-17 vs StpA2-17, not significant).

**Table 1.** Thermodynamic parameters for peptide partition to DSPC/DSPG membrane.

| | $K_x (\times 10^5)$ <sup>a</sup> | $\Delta G_x^0$ (kJ/mol) <sup>b</sup> |
| --- | --- | --- |
| **A2-17** | $2.7 \pm 0.19$ | $-31 \pm 0.17$ |
| **StpA2-17** | $2.4 \pm 0.14$ | $-31 \pm 0.15$ |
| **StchA2-17** | $110 \pm 16$ | $-40 \pm 0.34$ |

<sup>a</sup> The partition constant was measured using separation method<sup>39</sup> (see the Methods section for details).
<sup>b</sup> The standard Gibbs free energy was calculated according to $\Delta G_x^0 = -RT \ln K_x$ .

**Figure 4.**
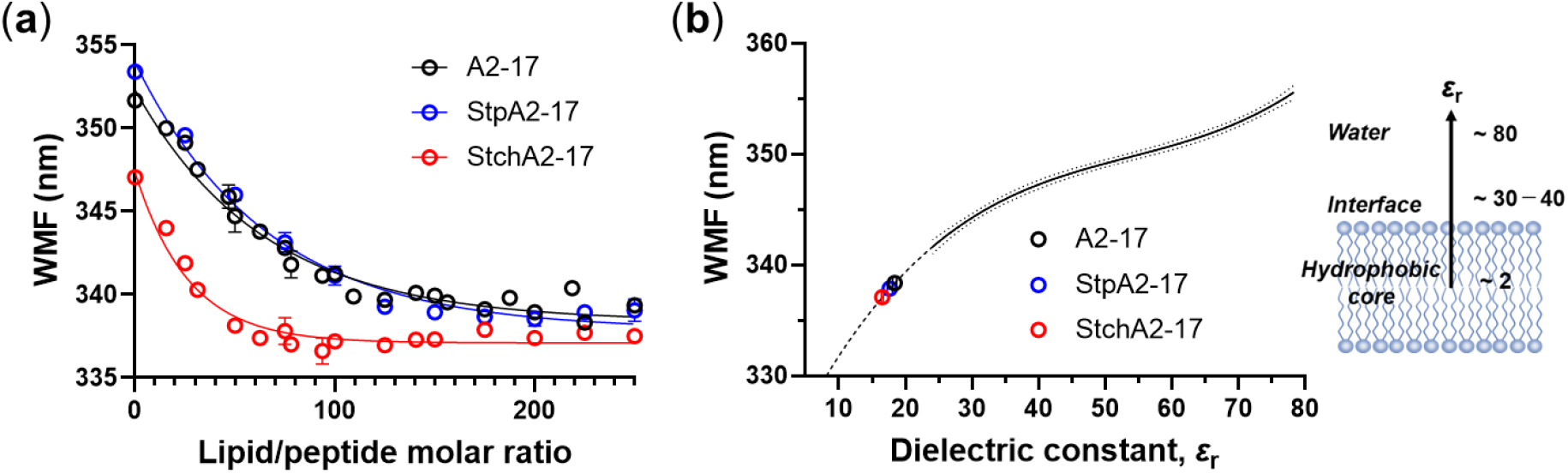
Lipid membrane binding of peptides. (a) Changes in WMF of Trp residue for A2-17, StpA2-17, and StchA2-17 as a function of molar ratio of total lipid of LUVs to peptide (lipid/peptide molar ratio changes from 0 to 250). One-phase decay curve fitting using the least squares method was applied to obtain the WMF at plateau. Solid lines represent the best fit. (b) The membrane location of the Trp residue in the peptide was estimated using the WMF profile of *N*-acetyl-L-tryptophanamide under different values of dielectric constant. The experimental data of *N*-acetyl-L-tryptophanamide was fitted with a third-order polynomial line that provided the best fit, and a 95% confidence band was included. The dashed line represents the extrapolation to a WMF value of 330, where the WMF values of A2-17, StpA2-17, and StchA2-17 at the plateau were fitted to the profile.

### Membrane perturbation and pore

In a previous study, we established an atomic force microscopy (AFM) method for measuring the mechanical membrane perturbation by peptides^18^. Using this method, we compared the membrane perturbation of A2-17 with that of StpA2-17 and StchA2-17 (Fig. 5). The spherical structure of LUVs was observed on the substrate in both the absence and presence of peptides (Fig. 5a), and the peptide-induced decrease in liposome stiffness was quantified as a membrane perturbation, as described in the Methods section. Despite its lower α-helicity and hydrophobicity, properties that generally enhance peptide partitioning to lipid membranes and membrane perturbation^18^, A2-17 caused a greater membrane perturbation than StpA2-17 and StchA2-17 (Fig. 5b). This result suggests that the structural flexibility of A2-17 contributes to increased membrane perturbation. We also compared the characteristics of peptide-induced pores in a planar dioleoylphosphatidylcholine (DOPC) bilayer membrane system using electrophysiological analyses (Fig. 6). Each current event caused by peptides can be classified into a “spike” or “long-lasting” signal based on the event duration, and the spike current signal (duration < 20 ms) is likely to be derived from the favorable trait of ARPs that can induce transient defects/pores of the lipid membrane to penetrate cells^18,30^. Figure 6a shows the typical current signals observed after peptide addition. Spike current signals were predominant for all the peptides: A2-17 (96.3%), StpA2-17 (99.6%), StchA2-17 (90.3%). The average frequency of spike current signals was in the order: A2-17 (0.2 s^-1^) > StpA2-17 (0.2 s^-1^) > StchA2-17 (0.005 s^-1^). In addition, the charge flux accompanied by spike current signal was much greater for A2-17 than that for StpA2-17 or StchA2-17 (Fig. 6b). These results indicate that A2-17 had the most favorable membrane perturbation mode for cell membrane penetration.

**Figure 5.**
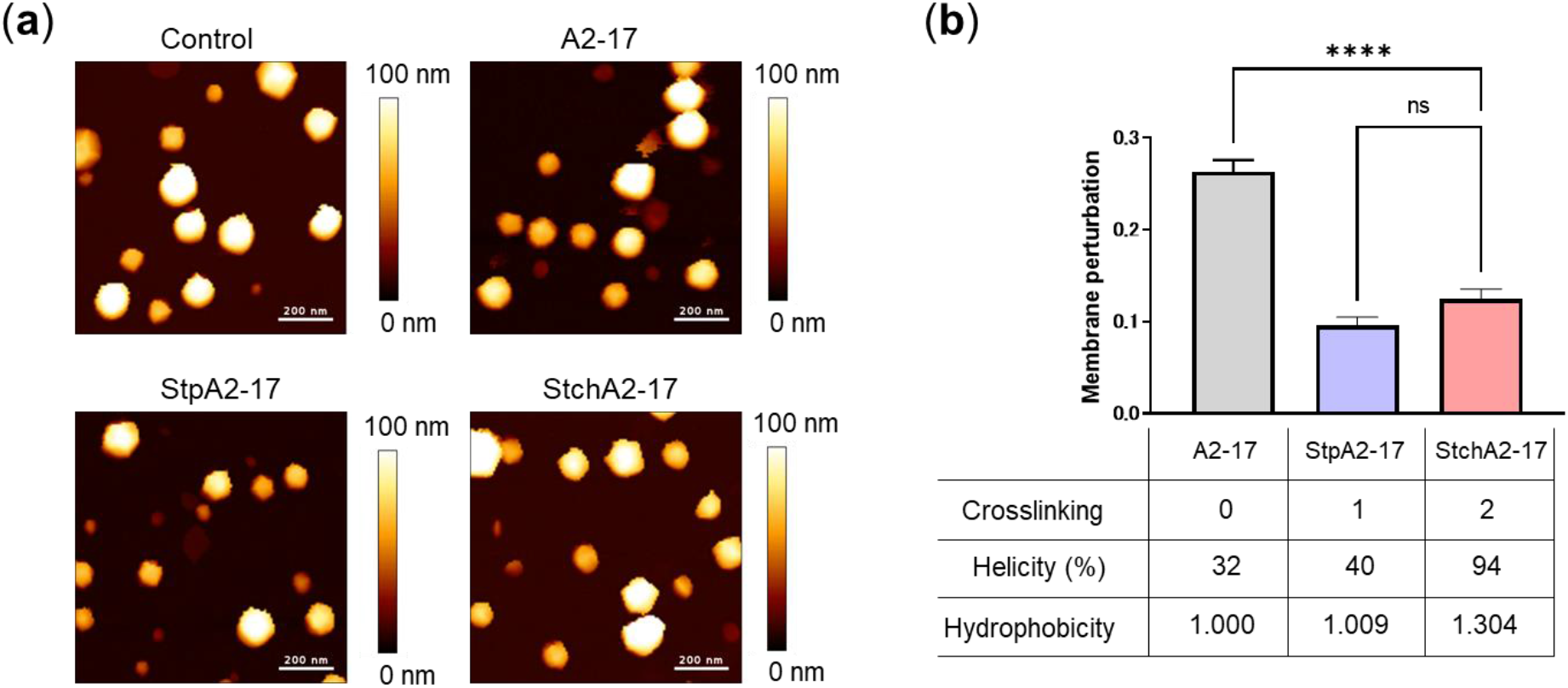
Peptide-induced membrane perturbation. (a) AFM images of LUVs in the absence (control) and presence of A2-17, StpA2-17, or StchA2-17. Scale bars represent 200 nm. (b) Membrane perturbations as defined by the rate of decrease in the stiffness of lipid vesicles. The table below the graph shows the crosslinking numbers (Crosslinking: stapled = 1; stitched = 2), α-helical contents bound to the LUVs from Fig. 1(b) (Helicity), and relative retention times (Hydrophobicity). \*\*\*\**p* < 0.0001; ns: not significant.

**Figure 6.**
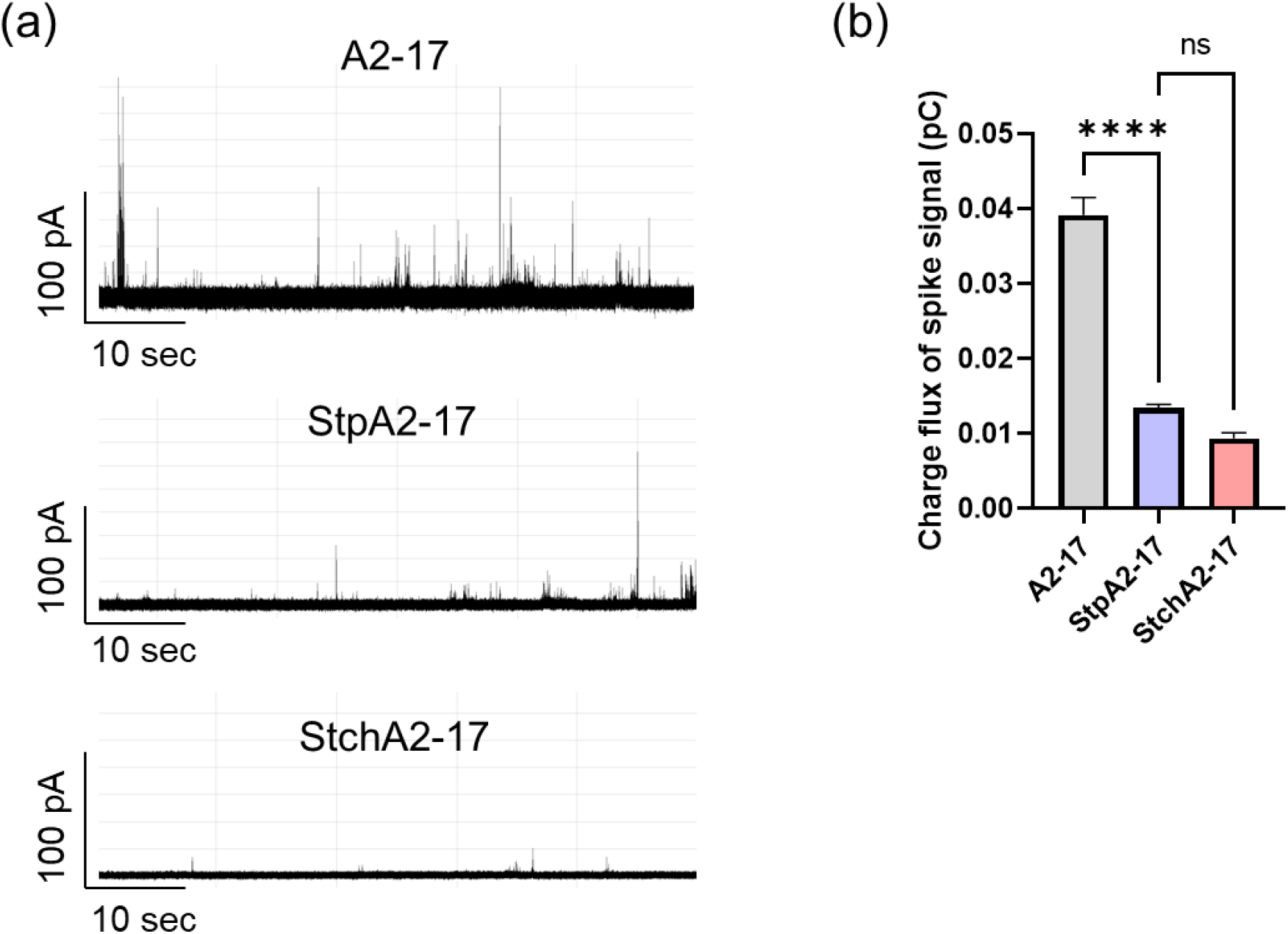
Channel current signals of A2-17, StpA2-17, and StchA2-17 after interaction with planar lipid membranes. (a) Typical current and time traces of A2-17, StpA2-17, and StchA2-17. (b) The charge flux of spike signals caused by the three peptides. \*\*\*\**p* < 0.0001; ns: not significant.

## Discussion

The conformation-restricting intramolecular linking has been known to enhance the cell penetration of peptides^20,24^. However, the present study showed that the introduction of hydrocarbon-stapling or hydrocarbon-stitching into structurally flexible A2-17, which changes conformation in response to different physicochemical environments, results in reduced cell membrane penetration ability.

It has been argued that structural flexibility is important for ARPs to adapt to hydrophilic and hydrophobic environments (aqueous phase and lipid membranes, respectively) so that they can penetrate cell membranes^6,10,22,25^. Although generalization is challenging owing to conflicting data^19,31,32^, stapling or stitching of amphiphathic ARPs could decrease entropy in the lipid-unbound state, thereby stabilizing the lipid-bound state and compromising their cell penetration capability. The reported enhanced cell penetration of stapled or stitched polyarginines^20,24^ may be derived from a moderate increase in amphipathicity or hydrophobicity by helical structures containing hydrophobic residues (S5, B5, and S8), which could improve their partitioning to lipid membranes. Our results suggest that for amphipathic ARPs, a stable helical structure with very high amphipathicity would make it difficult for the peptide to desorb from the plasma lipid membranes, resulting in poor cell penetration.

As previously discussed by Takechi-Haraya et al., the Born theory excludes the well-known solubility-diffusion mechanism of membrane penetration, where the permeating substance on one side first dissolves in the hydrophobic region of the membrane and then crosses the membrane to the other side according to the concentration gradient as the driving force^13^. Although StchA2-17 had more favorable energetics for membrane partitioning than A2-17 and StpA2-17, it consistently did not exhibit cell membrane penetration and remained on the cell membrane.

We found that the cell-penetrating A2-17 peptide softened the membrane mechanically, leading to transient membrane defects or pores^13,18^. The peptide-induced membrane perturbation, defined as a decrease in the mechanical rigidity (stiffness) of the interacting liposomes, was higher for A2-17 than for StpA2-17 and StchA2-17, as well as the increased charge flux associated with transient membrane pores caused by A2-17. StpA2-17 had lower cell penetration ability than A2-17 despite similarities in membrane insertion and partition energy; hence, it appears that A2-17 adopts favorable α-helical conformation with structural flexibility that causes the membrane perturbation required for efficient cell membrane penetration.

Chemical modifications that introduce conformational restrictions are generally advantageous for improving chemical stability, including peptidase resistance^21^. However, the present study highlights the need for careful consideration of such modifications to avoid potential loss of function in ARPs. Physicochemical characterization of the peptide in terms of secondary structure, peptide insertion and partitioning to lipid membranes, and membrane perturbation will be valuable for designing the optimal helical structures of ARPs. Additionally, conformational restriction due to steric hindrance of the side chain hydrophobic groups at the amino acid residue level reportedly controls the cell membrane penetration of cyclic peptides^23^. Therefore, more sophisticated tuning of a stapled or stitched helical structure with optimized amino acid residues may lead to the development of novel cell-penetrating peptides. One potential way to improve the cell membrane penetration ability of a hydrocarbon-stapled peptide would be to test various positions of the staple in the helical peptide. It has been shown that extending the hydrophobic face of an amphipathic helix by the placement of a staple between the hydrophilic and hydrophobic sides of the helix increases peptide’s cell penetration ability^31^.

In conclusion, A2-17 exhibited higher ability to penetrate cell membranes than StpA2-17, and StchA2-17 did not penetrate cells. StchA2-17 had much higher affinity for lipid membranes than A2-17 and StrpA2-17. Despite the similar degrees of membrane insertion for all peptides, A2-17 reduced the mechanical rigidity of liposomes to a greater extent than StpA2-17 and StchA2-17. Additionally, A2-17 induced a higher charge influx through transient membrane pores than StpA2-17 and StchA2-17. Our results indicate that the structural flexibility of A2-17 leads to a membrane perturbation mode that contributes to its greater cell membrane penetration ability, thus providing new insights for the future design of ARPs for drug delivery.

## Methods

### Materials

The peptide sequence of A2-17, StpA2-17, and StchA2-17 are shown in Fig. 1. The amino and carboxyl termini of each peptide were acetylated and amidated, respectively. For the FAM-labeled peptides, the amino terminus was labeled with FAM via a glycylglycine linker. A2-17 was purchased from the Peptide Institute, Inc. (Osaka, Japan). FAM-labeled A2-17, StpA2-17, and FAM-labeled StpA2-17 were purchased from GenScript Japan (Tokyo, Japan), and StchA2-17 and FAM-labeled StchA2-17 were purchased from WuXi AppTec (Shanghai, China). The purity of each peptide was certified to be >95% by the manufacturers. Stock solution concentrations of peptide and FAM-labeled peptide were prepared in water and determined by measuring absorbances of Trp at 280 nm (5500 M^-1^ cm^-1^)^33^ and 5-FAM at 494 nm (68,000 M^-1^ cm^-1^)^34^, respectively, using a NanoDrop One^c^ (Thermo Fisher Scientific, Waltham, MA, USA). DSPC, DSPG, and DOPC were purchased from Avanti Polar Lipids Ltd (Alabaster, AL, USA). L-Tryptophan and *N*-acetyl-L-tryptophanamide were purchased from Sigma-Aldrich (St. Louis, MO, USA).

### Preparation of lipid vesicles

LUVs were prepared as described by Takechi-Haraya et al^35^. Briefly, a dried lipid film of DSPC/DSPG (4:1 molar ratio) was hydrated with 10 mM Tris buffer containing 150 mM NaCl (pH 7.4) under mechanical agitation for 5 min at 60 °C. The resultant suspension was freeze-thawed five times using dry ice-methanol slush and a water bath of 60 °C. Thereafter, the suspension was extruded 21 times through a mini-extruder equipped with a 0.1-μm polycarbonate filter (Avanti Polar Lipids, Alabaster, AL, USA) at 70 °C.

### Analyses of peptide properties

To evaluate the secondary structure of peptides, far-UV CD spectra were recorded from 190 to 250 nm at 25 °C using a CD spectrometer (J-1100, JASCO, Tokyo, Japan) with a quartz cuvette of 1-mm path length. Peptide solutions (10 µM) in 10 mM Tris buffer containing 150 mM NaCl (pH 7.4) were subjected to CD measurements. This was in the absence as well as presence of LUVs (1 mM). Each CD spectrum of the peptide sample was corrected by subtracting the corresponding baseline for the same concentration of LUVs in Tris buffer. The α-helix content of peptide was determined from the mean residue ellipticity at 222 nm, as described by Scholtz et al^36^. CD measurements were performed three times for each group (*n* = 3). To evaluate the hydrophobicity of StpA2-17 or StchA2-17 relative to A2-17, their respective retention times were measured by reversed-phase high performance liquid chromatography using a Phenomenex Gemini 150 mm × 4.6 mm C18 110 Å 5 µm column (Phenomenex, Aschaffenburg, Germany) at 40 °C. The peptides were eluted using a linear gradient over 20 min at a flow rate of 1 mL/min, ranging from 5% to 65% acetonitrile/water containing 0.1% trifluoroacetic acid, with UV detection at 220 nm. The retention measurement was repeated three times and independently performed in triplicate (*n* = 9).

### Trp fluorescence measurements

Trp emission fluorescence spectra of 8 µM of A2-17, StpA2-17 or StchA2-17 in the absence and presence of LUVs at 25 °C in 10 mM Tris buffer containing 150 mM NaCl (pH 7.4) were record from 300 to 420 nm at an excitation wavelength of 290 nm using an F-7000 fluorescence spectrophotometer (Hitachi, Tokyo, Japan). To minimize the scattering artifacts from LUVs, with a polarizer orientation (Ex_pol_ = 90°, Em_pol_ = 0°), each Trp fluorescence spectrum of the peptide was corrected by subtracting the baseline for the same concentration of lipid vesicles in buffer solution, and further corrected for decreases in fluorescence intensity by using reference samples of 8 µM L-tryptophan with or without LUVs, as described by Ladokhin et al^27^. The WMF of each spectrum was determined by peak analysis using OriginPro software v.2023 (OriginLab Corporation, Northampton, MA, USA). In the same experimental condition, fluorescence spectra of *N*-acetyl-L-tryptophanamide (10 µM) in ethanol/water solvent was measured, and its dielectric constant-dependent WMF profile was obtained by the third order polynomial fitting to the data using the GraphPad prism software v.9.5.1 (GraphPad Software, La Jolla, CA). The dielectric constant of ethanol/water solvent was calculated from the equation: *ε*_r_ = 78.43*V*_water_ + 24.09*V*_ethanol_, where 78.43 and 24.09 are *ε*_r_ values of water^37^ and ethanol^38^ at 25 °C, respectively, and *V*_water_ and *V*_ethanol_ are volume fraction of aqueous buffer and ethanol, respectively. The standard Gibbs free energy of transfer from water to lipid membrane 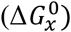 was calculated using the partition model^39^: 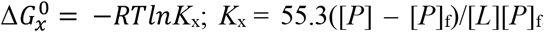, where *K*_x_ is the partition constant, [*P*] and [*L*] are the total molar concentrations of peptide and lipid, respectively, and [*P*]_f_ is the bulk molar concentration of peptide in water phase. For this analysis, at 25 °C in 10 mM Tris buffer containing 150 mM NaCl (pH 7.4), after 10 µM peptide was incubated for 30 min with or without LUVs (500 µM total lipid) in a 100 KDa MWCO Amicon Ultra^®^ tube (Merck Millipore, Billericia, MA, USA), the sample was centrifuged at 9000 × *g* for 15 min. For the resultant filtered solutions, Trp fluorescence intensities attributable to [*P*]_f_ and [*P*], which are samples with and without LUVs, respectively, were measured using a Synergy H1 plate reader (BioTek, Winooski, VT) at an excitation wavelength of 290 nm and a detection wavelength of 350 nm in flat-bottom 96-well plates. Experiments were conducted by preparing three to six samples from each group (*n* = 3–6).

### CLSM

LSM observation via the z-stack imaging mode was performed by a slight modification of our previous procedure^9^, using a confocal microscope (Nikon A1, Tokyo, Japan) with a 20× objective lens (CFI Plan Apo Lambda 20x, NA 0.75, WD 1.00 mm) at an excitation wavelength of 488 nm to visualize FAM-labeled peptides. Z-stack images were acquired from the periphery of the cells attached to the glass-bottom dish toward the opposite periphery by scanning every 0.78 μm. The detection pinhole diameter was set to be 1.0 times of the diameter of the Airy Disk, resulting in an optical section thickness of < 2.1μm. HeLa cells (2 × 10^5^ cells) were plated in a 35-mm glass-bottomed dish coated with poly-L-lysine (Matsunami Glass Ind. Ltd., Osaka, Japan) and were incubated in Dulbecco’s modified Eagle medium (DMEM; Thermo Fisher Scientific) supplemented with 10 % fetal bovine serum (Sigma-Aldrich). After incubation for 24 h (37 °C, 5% CO_2_), the cells were incubated with 1 µM FAM-labeled peptide for 30 min at 4 °C in DMEM. After incubation, the cells were washed three times with phosphate-buffered saline on ice and stored in the phosphate-buffered saline, followed by confocal microscopy. The nuclei of the cells were counterstained with Cellstain^®^ Hoechst 33342 solution (DOJINDO LABORATORIES, Kumamoto, Japan) following the manufacturer’s instructions and visualized at an excitation wavelength of 405 nm. Throughout the image acquisition, the laser intensity, photomultiplier detector sensitivity, and pinhole aperture values were kept constant. ImageJ Fiji software^40^ was used to analyze the FAM fluorescence intensity of the nucleus region by subtracting the background intensity outside the cells, and the cell penetration of peptides was compared. Fluorescence analysis was performed on more than 40 cells (*n* = 40–125).

### Channel current analysis for membrane penetration of peptides

Electrophysiological measurements were performed using a MECA 16 TC chip with an Orbit 16 TC device (Nanion Technologies GmbH, Munich, Germany) according to the manufacturer’s instructions. The MECA 16 TC chamber was filled with 200 µL of buffer solution (150 mM KCl, 10 mM MOPS, pH 7.0); 1 µL of DOPC (lipid/n-decan, 10 mg/mL) solution was added to the chamber, and planar lipid bilayers were automatically prepared using the Orbit 16TC stirrer. The peptide was then dissolved in the chamber at 10 µM (a lipid/peptide molar ratio of ∼ 6 in the chamber system), and 100 mV of voltage was applied. The channel current was monitored using the Orbit 16 TC Elements Data Reader 4 v.1.0.22. The channel current signals were detected using a 4-kHz low-pass filter at a sampling frequency of 20 kHz. Analysis of the channel signals was performed using pCLAMP ver. 10.7 (Molecular Devices, CA, USA). The data obtained in this measurement are *n* (number of current signals): 2081 > *n* > 311; *N* (number of experiments) > 3.

### AFM

To measure the membrane perturbation of peptides, we performed AFM at 25 ± 1 °C in 10 mM Tris buffer containing 150 mM NaCl (pH 7.4) using a BioLever mini cantilever (BL-AC40TS, Olympus Co., Tokyo, Japan) via the QI mode of a JPK Nanowizard Ultra Speed microscope equipped with the Data Processing JPK software v.6.0 (JPK Instruments AG, Berlin, Germany) according to our previous procedure^18^. Briefly, 200 μL of DSPC/DSPG-LUVs (50 μM of total lipids) in Tris buffer solution was incubated on an aminopropyl-modified mica substrate for 20 min, and an additional 1.4 mL of Tris buffer solution with or without the peptide was added, followed by AFM measurement. For the peptide samples, the final peptide/lipid molar ratio was 1. The AFM images were recorded at a resolution of < 8 nm/pixel. To obtain the lipid vesicle stiffness, a linear fit was performed over the linear region of the force-deformation curve at the center of a lipid vesicle using JPK Software. Peptide-induced membrane perturbation was defined as a decrease in lipid vesicle stiffness using (*S*_control_ – *S*)/*S*_control_, where *S*_control_ and *S* are the stiffnesses of control lipid vesicles and lipid vesicles treated with peptide, respectively^18^. Using the average stiffness for *S*_control_, we evaluated the membrane perturbation of lipid vesicles with peptides. Small lipid vesicles exhibit increased stiffness due to their relatively high membrane curvature^41^. In this study, we analyzed the stiffness data of liposomes with a height (*h*) described by the equation 10*x* + 10 > *h* ≧ 10*x*, where *x* ranged from 5 to 14. Membrane perturbations were then obtained for each *x* group and averaged across all perturbations. The experiment was repeated three times and AFM analysis was performed on a total of 176 to 384 liposomes (*n* = 176–384).

### Statistical analysis

The results are presented as mean ± standard error. Statistical analyses were performed using GraphPad prism version 9.5.1. Differences between groups were analyzed using a one-way ANOVA with Tukey’s multiple comparison test. The results were considered statistically significant at a *p*-value < 0.05.

## Acknowledgements

The authors express their deepest appreciation to Dr. Yosuke Demizu (National Institute of Health Sciences) for supporting the CD measurements. The authors thank Editage [https://www.editage.com/] for English language editing and manuscript review. This study was partially supported by JSPS KAKENHI Grant Numbers JP20K15982 and 23K06092 (to Y.T-H.) from Japan Society for the Promotion of Science, and by AMED under Grant Number JP23mk0101197h0203 (to K.I.).

## Author Contributions

Y.T-H.: conception and design; acquisition, analysis, and interpretation of data; drafting and revising of manuscript. T.O.: interpretation of data; revising of manuscript. A.U.: acquisition of data; revising of manuscript. K.N.: interpretation of data; revising of manuscript. K.U.: interpretation of data; revising of manuscript. Y.A.: interpretation of data; revising of manuscript. R.K.: interpretation of data; revising of manuscript. M.I.K.: conception; interpretation of data; revising of manuscript. M.R.: conception; interpretation of data; revising of manuscript. A.T.R.: conception; interpretation of data; revising of manuscript. Y.S.: supervision, interpretation of data; revising of manuscript. K.I.: supervision, interpretation of data; revising of manuscript. H.S.: conception and design; interpretation of data; revising of manuscript. All authors reviewed the results and approved the final version of the manuscript.

